# A Cell Cycle-aware Network for Data Integration and Label Transferring of Single-cell RNA-seq and ATAC-seq

**DOI:** 10.1101/2024.01.31.578213

**Authors:** Jiajia Liu, Jian Ma, Jianguo Wen, Xiaobo Zhou

## Abstract

In recent years, the integration of single-cell multi-omics data has provided a more comprehensive understanding of cell functions and internal regulatory mechanisms from a non-single omics perspective, but it still suffers many challenges, such as omics-variance, sparsity, cell heterogeneity and confounding factors. As we know, cell cycle is regarded as a confounder when analyzing other factors in single-cell RNA-seq data, but it’s not clear how it will work on the integrated single-cell multi-omics data. Here, we developed a Cell Cycle-Aware Network (CCAN) to remove cell cycle effects from the integrated single-cell multi-omics data while keeping the cell type-specific variations. This is the first computational model to study the cell-cycle effects in the integration of single-cell multi-omics data. Validations on several benchmark datasets show the out-standing performance of CCAN in a variety of downstream analyses and applications, including removing cell cycle effects and batch effects of scRNA-seq datasets from different protocols, integrating paired and unpaired scRNA-seq and scATAC-seq data, accurately transferring cell type labels from scRNA-seq to scATAC-seq data, and characterizing the differentiation process from hematopoietic stem cells to different lineages in the integration of differentiation data.

## INTRODUCTION

In recent years, advances in single-cell RNA sequencing (scRNA-seq) technology have enabled us to generate high-throughput gene expression data through different sequencing methods at single-cell resolution [1–4]. The evolution of these technologies has significantly expanded the adoption of single-cell RNA sequencing across diverse fields, furnishing a more comprehensive and profound perspective to the comprehension of cell heterogeneity and functions [3, 5–7]. There exists a multitude of methods for generating scRNA-seq data, with over a dozen most commonly used scRNA-seq protocols accessible [8]. These technologies generate scRNA-seq data derived from distinct experiments, encompassing variations in capture timing, handlers, reagent batches, equipment, and even technological platforms [8–11]. These inherent dissimilarities engender batch effects within scRNA-seq data [6, 12], which become the priority and grand challenge in single-cell RNA-seq data analysis [13].

In addition, the emergence of single-cell multi-omics technologies has enabled insights into complex cellular microenvironments and biological processes, offering many exciting biological opportunities from perspectives other than transcriptomics, such as genomics, epigenomics, proteomics, metabolomics and spatial transcriptomics [14–16], etc. Particularly, single-cell Assay for Transposase-Accessible Chromatin using sequencing (scATAC-seq) is an epigenomic profiling technology for studying the chromatin accessibility of individual cells [17–19]. It provides the ability to examine the openness of chromatin regions in the nucleus at the single-cell level [17], which is unavailable in single-cell RNA-sequencing data. This enhances our comprehension of the epigenetic state, cell-type heterogeneity and cell state [20, 21]. However, scATAC-seq data has extreme sparsity than scRNA-seq data [18, 22, 23], which also increases the difficulty of analysis based on scATAC-seq data. Meanwhile, cell type annotation in scATAC-seq data is challenging due to lack of specifically designed tools and use of unintuitive cis- and trans-regulatory elements in single cell ATAC-seq data [24]. In recent years, advanced technologies [25–28] make it possible to simultaneously characterize gene expression and chromatin accessibility in the same cell, which we called the generated data paired data. These techniques provide tools for the integrated analysis of scRNA-seq and scATAC-seq data, which can apply the information obtained from the large amount of annotated scRNA-seq data for the cell type annotation of scATAC-seq data [29]. However, single-omics data are more readily available than paired scRNA-seq and scATAC-seq data. That is, we can generate scRNA-seq and scATAC-seq data of different single-cell experimental samples separately, but they are from the same organ or tissue [2], which we refer to as unpaired data in this study. Therefore, in cases where paired data are not abundant, integrating these unpaired data is a better option for researchers to conduct broader studies. However, unpaired data frequently introduce complexity to subsequent analyses due to differences in features and sparsity level, so it is important to develop novel data integration methods that can be applied to unpaired scRNA-seq and scATAC-seq data.

In summary, the current integration of scRNA-seq and scATAC-seq data can be divided into three types [30]: (1) intra-modality integration, the integration of the same omics data (scRNA-seq data) measured from different cells and different experiments; (2) paired inter-modality integration, that is, the integration of scRNA-seq and scATAC-seq data measured from the same cell; and (3) unpaired inter-modality integration, that is, the integration of scRNA-seq and scATAC-seq data generated from different cells, samples or experiments. Corresponding integration methods have also been developed for different integration types: (1) methods for intra-modality integration [31–34] employ dimensionality reduction algorithms to reduce the complexity of the data and identify the common biological signal across datasets to align cells or cell populations to integrate scRNA-seq datasets from different sources or experiments, but most of them struggle with excessive data scale, run time or resource requirements. (2) methods for paired inter-modality integration apply matrix factorization [35], weighted nearest neighbor algorithm [36] and neural networks [37, 38] to integrate scRNA-seq and scATAC-seq data measured within a cell and to obtain a joint definition of cellular state. These methods are specially designed for paired data, making their application to other unpaired data challenging. (3) unpaired data not only have different features, but also frequently exhibit significant variations in cell count, thus giving rise to a distinct category of integration methods for unpaired data. Methods for unpaired inter-modality integration focus on finding solutions of the manifold learning and cell alignment in the embedding space using neural networks [39–41]. There is also a non-neural network approach that uses the non-negative matrix factorization approach and online learning algorithm to incorporate new data without recalculating from scratch [42]. However, despite aforementioned approaches, most of them are specialized to address specific challenges within particular integration type of single-cell data. Currently, there is a lack of a comprehensive approach capable of simultaneously addressing all three types of integration issues outlined above. The single-cell data integration method to be developed needs to consider the sparsity, data scale, feature difference, high dimensionality and other inherent disparities of scRNA-seq and scATAC-seq data. In addition, the cell cycle is often considered a confounding factor in the study of cell population and cell heterogeneity based on single-cell RNA-sequencing data [43–47]. While how it will work in the integrated analysis of scRNA-seq and scATAC-seq data is still unknown.

To address such concerns, we developed an advanced Cell Cycle-Aware Network (CCAN) with the aim of extracting intrinsic biological signals masked by context-specific patterns (i.e., cell type-specific heterogeneity) and confounding factors (i.e. cell cycle effects, batch effects, and noise). Notably, CCAN can integrate single-cell multi-omics data and remove cell cycle effects from the integrated data while maintaining heterogeneity between cell types. CCAN is based on a domain separation network, adding a periodic activation function to the private decoder to simulate the dynamic process of the cell cycle, and projecting single-cell data from different platforms or modalities into a common low-dimensional space through shared projection. The distribution constraint function and the class alignment loss function are added to the shared embedding space to make the distribution of different data as similar as possible and the difference between different types of data to be maximized. In addition to single-cell data integration, CCAN enables cell type prediction of scATAC-seq data via transferring the cell type annotation information of scRNA-seq data to scATAC-seq data. Validations based on multiple sets of data prove that CCAN can not only eliminate the batch effect between scRNA-seq data from different platforms, but also integrate paired and unpaired scRNA-seq data and scATAC-seq data well in the embedding space. Integration of unpaired data enables accurate cell type prediction for scATAC-seq data. Furthermore, CCAN can maintain cell differentiation trajectories when integrating single-cell differentiation data.

## RESULTS

### Overview of CCAN approach

As illustrated in **Fig. 1**, CCAN is a self-supervised approach using the labeled transcriptomic profile of scRNA-seq data (source domain) and unlabeled profile from same/different omics data (target domain), such as gene expression of scRNA-seq data and chromatin accessibility of scATAC-seq data. CCAN uses a domain separation network (DSN) to integrate data from source and target domains and transfer the annotations from source domain to target domain. Shared encoder and private encoders in DSN are three-layer perceptrons to learn noncircular and circular embeddings of both domains, separately. The shared embedding function projects a high-dimensional profile of each cell to a low-dimensional vector, which distinguishes biological meaningful signals from circular confounding factors (private embedding) and transforms the embeddings of cells from different domains into a similar distribution. In the decoder, we use sine and cosine as the activation functions specific for private embeddings, followed by a two-layer perceptron performing noncircular transformations mapping the embedded data to the original space. The training of CCAN has four main steps: (1) pretraining of the cell cycle-aware domain separation network; (2) label transferring from source domain to target domain; (3) refining CCAN by introducing a cluster alignment loss and (4) finalization and applications of CCAN model. We assessed CCAN using several real single-cell datasets, including two scRNA-seq datasets from different protocols [33], paired and unpaired scRNA-seq and scATAC-seq datasets [27, 48] and single-cell differentiated datasets from different modalities [49]. Evaluation results indicate that CCAN is a versatile method that can be used for multi-tasks, including single-cell multi-omics data integration, batch effect removal, cell cycle effects removal, label transferring and cell type prediction (**Fig. 1**). CCAN is an effective method and competitive in various applications compared with other existing methods.

**Fig. 1.**
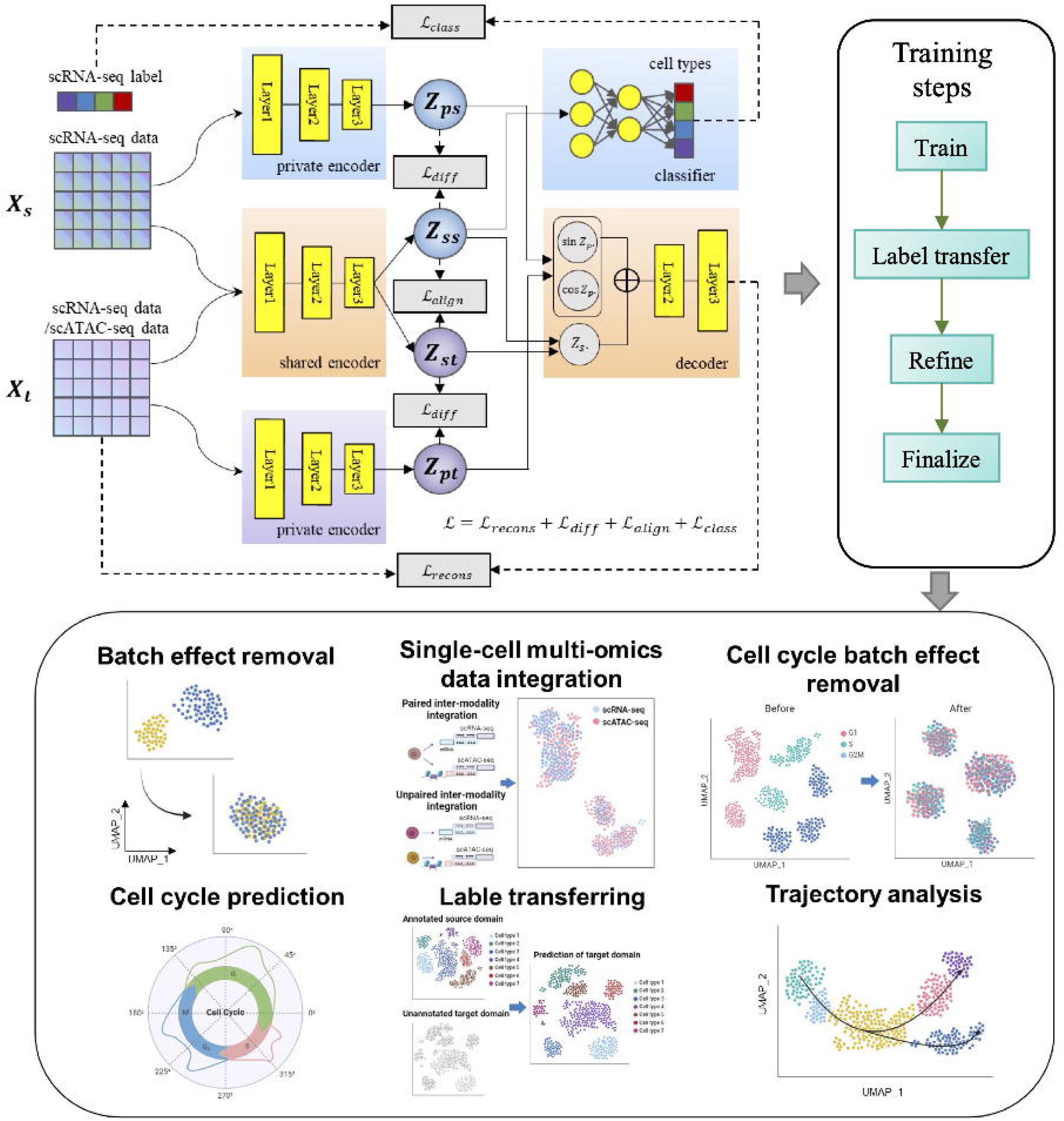
Overview of CCAN approach. CCAN is based on a domain separation network, taking labeled scRNA-seq data as source domain and unlabeled scRNA-seq or scATAC-seq data as target domain. Both source and target domain are fed into a shared encoder and two private encoders. The shared encoder learns the common information from both domains and employs an alignment loss to constrain the distribution of source and target domain to be similar in the shared embedding space. Private encoders are specific to source or target domain to extract periodic information of the cell cycle effect. An orthogonal difference loss between the shared embedding and private embedding enforces features learned by the shared encoder and the private encoder to be as different as possible in the low-dimensional space. Both source and target domain have the same structure of decoder by contacting the shared embedding and private embedding to reconstruct the original data. A supervised classifier is trained on the labeled scRNA-seq data and can be used to predict the label of target domain. CCAN has four main training steps and can be applied to multi-tasks, including batch effect removal, single-cell multi-omics data integration, cell cycle effect removal, cell cycle prediction, label transferring and trajectory analysis, etc.

### Batch effect removal of scRNA-seq datasets from different protocols

The emergence of advanced technologies enabled comprehensive transcriptional characterization of cell-type heterogeneity across a variety of biological and clinical conditions, integrating these scRNA-seq datasets from different protocols while maintaining cell-type heterogeneity has become very challenge. To benchmark the performance of CCAN against other existing methods for scRNA-seq data integration, we applied two scRNA-seq datasets of human Peripheral Blood Mononuclear Cells (PBMC), each assayed on the Chromium 10X platform but prepared with different protocols: 3’ end v1 and 3’ end v2 chemistries [33]. We denoted them as pbmc_6k and pbmc_8k respectively in this study. We first clustered the two scRNA-seq datasets separately using the Louvain clustering algorithm [50], resulting 10 clusters of the pbmc_6k data and 13 clusters of the pbmc_8k data (**Fig. 2a**). Then we used canonical cell type marker genes to annotate PBMC clusters before integration (**Fig. 2b, Supplementary Fig S1&S2**). The annotation resulted six major cell populations (**Fig. 2c**): monocytes (CD14+/FCGR3A+/MS4A7+/LYZ+), dendritic cells (FCER1A+), B cells (MS4A1+), T cells (CD3D+), megakaryocytes (PPBP+) and natural killer (NK) cells (CD3D-/GNLY+) [36, 51]. These marker genes were significantly expressed in the corresponding annotated cell types (**Fig. 2d**). We compared CCAN with six existing methods for scRNA-seq integration, including Harmony [33], Seurat V4 [36], online iNMF [42], Conos [31], Scanorama [32] and BBKNN [34]. As illustrated in **Fig. 2e**, the two PBMC data were completely separated before integration, which showed the confident existence of batch effect. These batch effects significantly impeded the accurate classification of cell types. Evidently, the same cell types were subdivided into two distinct categories prior to integration, including B cells, T cells, and monocytes. CCAN outperformed other methods by accomplishing a perfect integration, effectively eliminating dataset-specific variations and mixing all cells within each cluster in the projected space. The projection distribution on the two coordinate axes of UMAP (Uniform Manifold Approximation and Projection for Dimension Reduction) [52] visualization also illustrated that the distributions of pbmc_6k and pbmc_8k exhibited near-complete overlap, further affirming the successful integration of PBMCs using CCAN. To quantitatively measure the performance of CCAN, we calculated kBET (k-nearest neighbor Batch Effect Test) [53], a metric used to assess the batch effects in single-cell RNA sequencing data by comparing the distribution of nearest neighbors between cells from different batches. Compared to other methods, CCAN had the lowest kBET score when choosing different numbers of nearest neighbors (**Fig. 2f**), indicating that cells from different batches had more similar nearest neighbors after integration using CCAN. What’s more, a perfect integration not only required no significant variability between the two scRNA-seq data, but also needed to enhance the differences between cell types, which was valuable for subsequent downstream analyses. We applied k-means clustering to the integrated data, and compared the clustering results with the annotated cell types. By calculating the three clustering evaluation metrics of Rand Index (RI), Adjusted Rand Index (ARI), and Normalized Mutual Information (NMI), we observed that the results of CCAN were slightly inferior to those of BBKNN and Seurat V4, but always higher than Harmony, online iNMF, Conos and Scanorama (**Fig. 2g**). However, considering the performance of both data mixing and cell type separation, CCAN was still a better choice for integration that had less impacted by batch effects and was more reliable for downstream analyses.

**Fig. 2.**
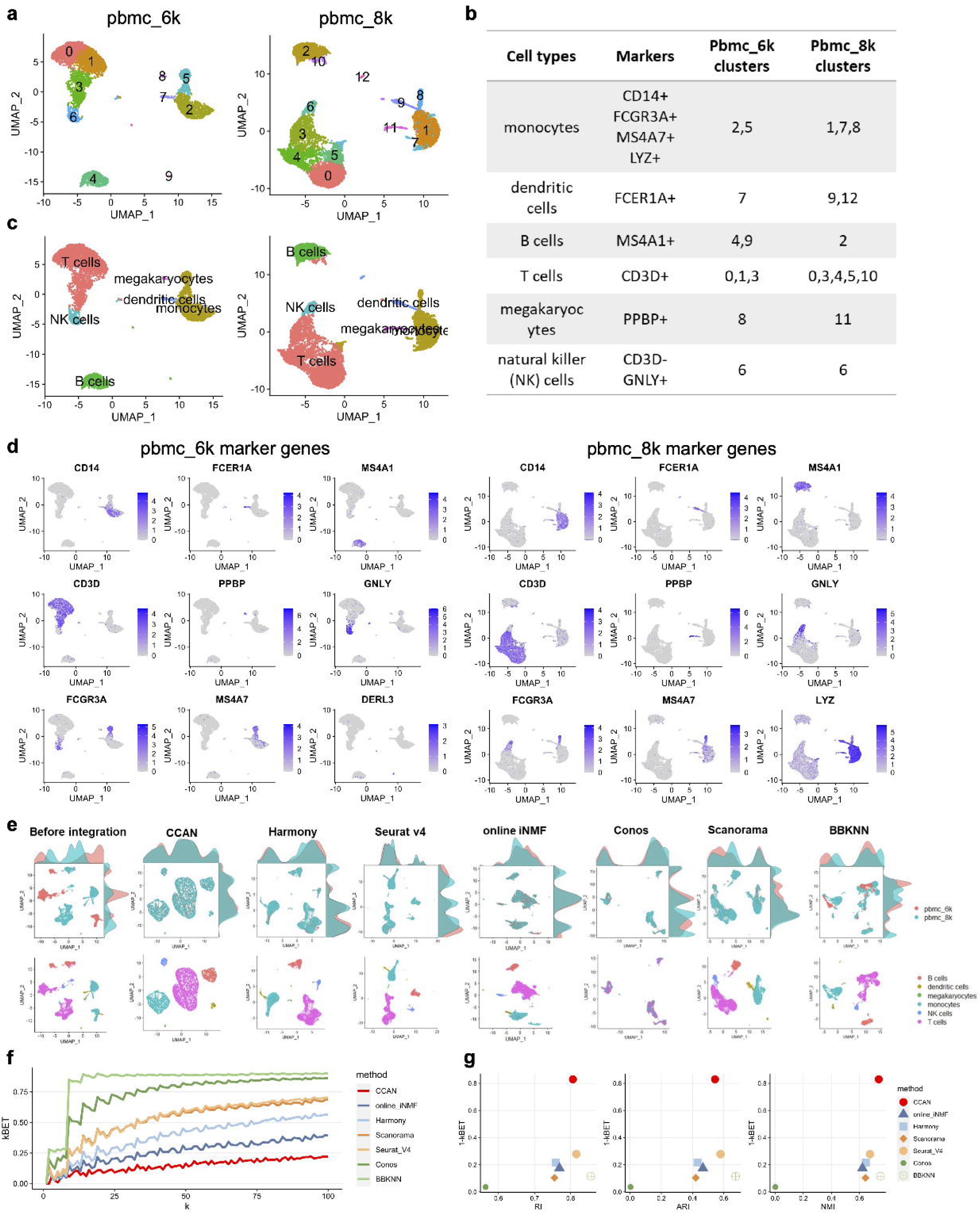
Annotation of PBMCs and batch effect removal of scRNA-seq datasets from different protocols. **a,**UMAP visualization of two single-cell PBMC data from different protocols. Different colors represent different clusters using Louvain clustering method. **b,** Annotation of different clusters of two PBMC data. Different cell clusters were annotated into six major cell types using marker genes. **c,** UMAP visualization of two single-cell PBMC data from different protocols with annotated cell types. **d,** UMAP visualization of marker expression in pbmc_6k and pbmc_8k data, respectively. **e,** UMAP visualization of scRNA-seq datasets before integration and after integration using CCAN, Harmony, Seurat V4, online iNMF, Conos, Scanorama and BBKNN, respectively. The upper panel was colored by different data batches and the bottom panel colored by annotated cell types. **f,** line plot of kBET scores of integrated data using different methods when choosing different numbers of nearest neighbors. k is neighborhood size. **g,** scatter plot of kBET score and three clustering metrics including RI, ARI and NMI of different methods. Different methods were represented using different colors and different shapes. RI, Rand Index; ARI, Adjusted Rand Index; NMI, Normalized Mutual Information.

### Cell cycle effect removal of the integrated data

In addition to the impact of data batches on cell type heterogeneity, cell cycle is often seen as a confounding factor when studying differences between cell types. Before integration, we meticulously selected a subset of variable genes from the PBMCs and proceeded to conduct Principal Component Analysis (PCA) based on these selected genes. Notably, some genes associated with the cell cycle prominently featured among the top 10 principal components. For instance, in the pbmc_6k dataset, genes such as *TYMS*, *RRM2*, *BIRC5*, *PCNA*, and *HMGB2* displayed significant associations with principal components 7, 8, 9 and 10 (PC_7, PC_8, PC_9 and PC_10). Similarly, within the pbmc_8k dataset, genes *TYMS*, *BIRC5*, and *MKI67* were prominent in principal component 10 (PC_10) (**Fig. 3a**). Despite their minimal or low expression in the majority of cells, several cell cycle-related genes (such as *PCNA* and *HMGB2* in pbmc_6k and *BIRC5* in pbmc_8k) exhibited normal expression patterns, thereby corroborating the presence of cell cycle effects. Inspired by Cyclum [45], we developed a cell-cycle aware module in CCAN to remove cell cycle effects in the integration of scRNA-seq data. CCAN employed a distinct sinusoidal component in the private autoencoder to effectively capture the circular trajectory in the high-dimensional gene expression space. The private embedding space was formed by single cells sampled at various stages of a periodic process. In essence, the private encoder in CCAN was dedicated to pinpointing an optimal cell embedding within this circular space, which we denoted as cell cycle pseudotime. We compared the performance of CCAN with other existing cell cycle effect removal methods, including Seurat, Cyclum and ccRemover. Before integration, we used the cell cycle marker genes in Seurat [36] to identify the cell cycle phases of cells in the two scRNA-seq datasets, and used the identified cell cycle as a benchmark label to evaluate the effectiveness of cell cycle effects removal. UMAP visualization of the integrated data after cell cycle effects removal showed that integration using CCAN were unable to clearly distinguish the three annotated cell cycle phases, as did Seurat, Cyclum and ccRemover. However, the advantage of CCAN over other methods was that only CCAN did not introduce extra noise to the downstream analysis based on cell types after removing the cell cycle effects, which was reflected in the fact that CCAN can accurately distinguish six different cell types after removing the cell cycle effects (**Fig. 3b)**. This also demonstrated the effectiveness and reliability of CCAN in removing cell cycle effects.

**Fig. 3.**
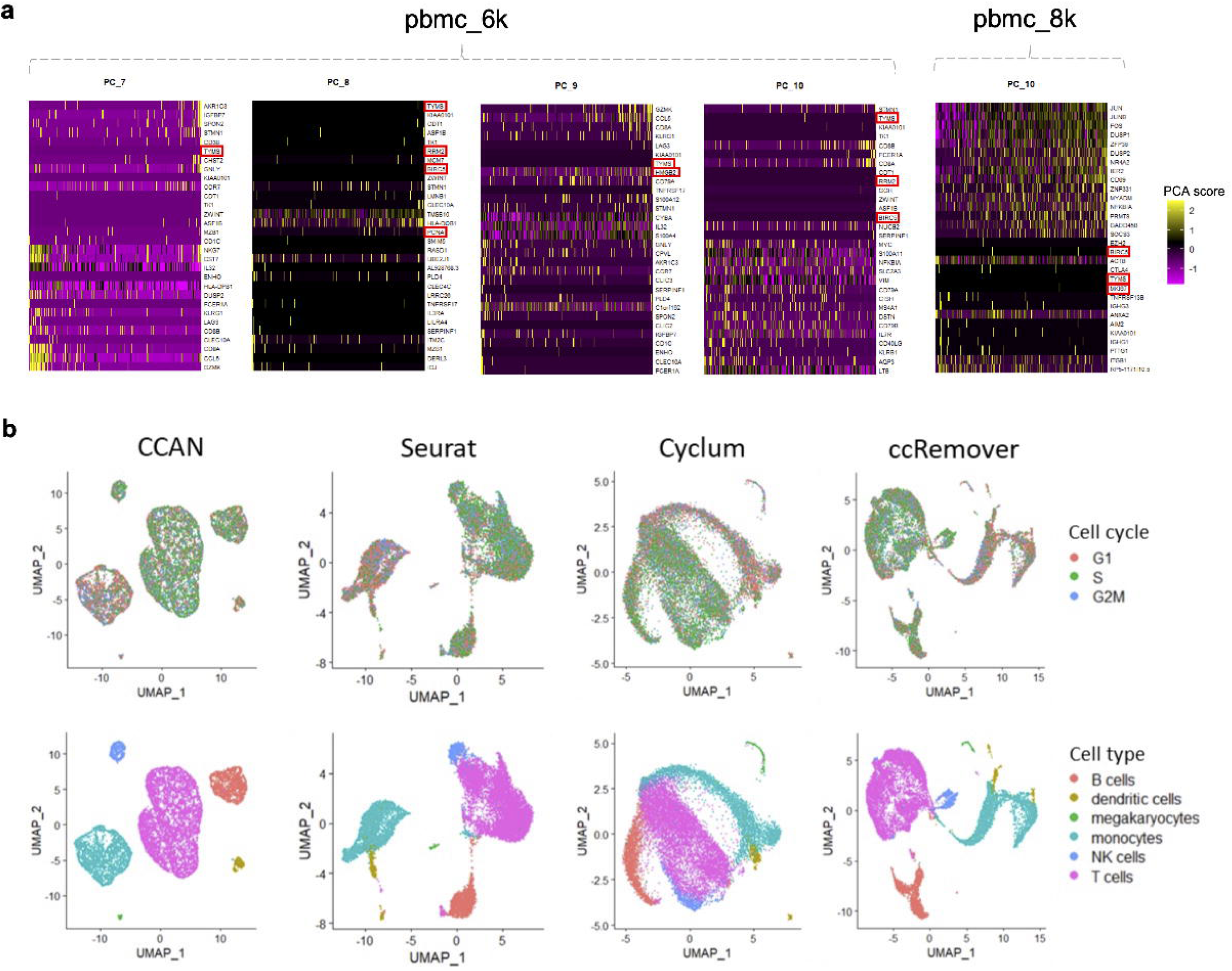
Cell cycle effect removal of integrated single-cell PBMC data. **a,** Heatmap of variable genes in principle components of PBMCs in two scRNA-seq data. Only the principal components related to cell cycle genes among the top ten principal components are shown. Cell-cycle genes are marked in red boxes. **b,** UMAP visualization of integrated PBMC data after removal of cell cycle effects using CCAN, Seurat, Cyclum and ccRemover. Top panels are colored by annotated cell cycle and bottom panels are colored by annotated cell types.

### Integration of joint profiling of scRNA-seq data and scATAC-seq data

Advanced technologies make it possible to simultaneously measure gene expression and chromatin accessibility in the same cell, such as SNARE-seq [27]. We applied CCAN to the paired single cell dataset, including joint profiling of scRNA-seq data and scATAC-seq data from adult mouse brain using SNARE-seq technology [27]. We compared the performance of CCAN with eight existing integration methods, containing scMVP [37], scAI [35], scMVAE [38], Seurat v4 [36], scJoint [40], GLUE [39], SCALEX [41] and online_iNMF [42]. Among these methods, scMVP, scAI and scMVAE are specifically designed only for integrating paired data, so their generation of the integrated data is based on concatenation of features in the latent dimension (**Supplementary Fig S3**). While Seurat v4, scJoint, GlUE, SCALEX and online_iNMF have the capability to integrate both paired and unpaired datasets. Their integration strategy for paired datasets in the comparison is to treat different modalities as two datasets from different experiments and integrate them together, so their integration is based on concatenation of cells (**Supplementary Fig S3**). The paired datasets are measured in the same cells, which provides groundtruth labels that allows CCAN to integrate without predicting the pseudo label of the scATAC-seq data first (**see MATERIALS AND METHODS**). In addition, CCAN’s integration for paired data is also the concatenation of learned features of scRNA-seq and scATAC-seq data in the latent space. UMAP visualization of the integrated data shows the excellent integration ability of CCAN, which enables the integrated data to distinguish 13 different cell types better than other methods. Following CCAN, GLUE exhibits the second-best performance. Notably, CCAN’s integrated data showcases a tighter aggregation of cells of the same type, with cells of different types are more dispersed. Although scAI, scMVAE, Seurat V4 and SCALEX can also clearly distinguish the large clusters such as L2 /3T, L4, L5CT, and L6IT, different cell types are not far away from each other (**Fig. 4a**). The results from three clustering evaluation metrics, including Rand Index (RI), Adjusted Rand Index (ARI), and Normalized Mutual Information (NMI), computed based on k-means clusters and ground truth, further corroborate that CCAN outperforms other methods in both integrating cells and accurately separating different cell types (**Fig. 4b**).

**Fig. 4.**
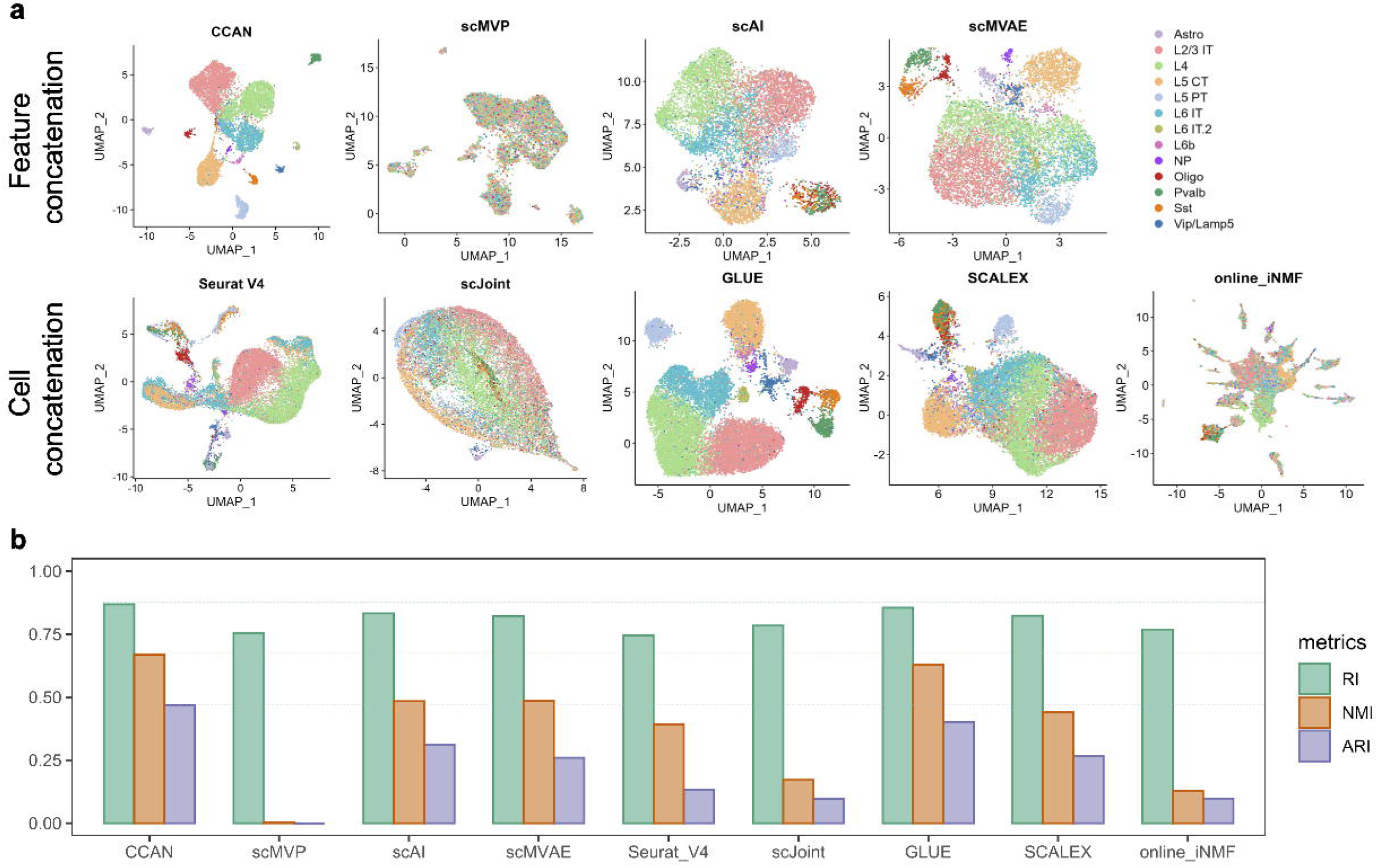
Data integration of paired datasets of scRNA-seq data and scATAC-seq data from adult mouse brain using SNARE-seq technology. **a,** UMAP visualization of integrated SNARE-seq data using CCAN, scMVP, scAI, scMVAE, Seurat v4, scJoint, GLUE, SCALEX and online_iNMF, respectively. Different colors represent different cell types annotated by Seurat. **b,** bar plot of three clustering metrics including RI, ARI and NMI of different methods. Different metrics were represented using different colors. RI, Rand Index; ARI, Adjusted Rand Index; NMI, Normalized Mutual Information.

### Integration of unpaired datasets and label transferring from scRNA-seq data to scATAC-seq data

Although more and more technologies are able to measure gene expression and chromatin accessibility in the same cell, there are still unpaired datasets from the same tissue. There is no correspondence between cells in scRNA-seq data and cells in scATAC-seq data. The integration of unpaired datasets maps both modalities to a common space, which allows tools and analyses designed for scRNA-seq data have the potential to be applied to scATAC-seq data. The variations between different modalities is different from batch effects between scRNA-seq datasets from different protocols, since scATAC-seq data characterize the chromatin accessibility instead of transcriptome profile and have more sparsity than scRNA-seq data. Even though integration of unpaired datasets from different modalities has many challenges, CCAN still has excellent performance compared with other existing methods (**Fig. 5a**). We assessed the performance of CCAN using scRNA-seq data from CITE-seq and scATAC-seq data from ASAP-seq. The CITE-seq generates filtered scRNA-seq data based on the condition of mitochondrial reads greater than 10%, number of expressed genes fewer than 500 and total number of UMI fewer than 1,000. For the scATAC-seq data from ASAP-seq, we filtered out cells with a number of peaks more than 100,000 and calculated the gene activity matrices for scATAC-seq data using Signac [54]. After that, 17,668 overlapped genes are selected as the input features of CCAN. In comparison to the state before integration, CCAN effectively integrates data from two distinct modalities, leading to the clear identification of seven cell types in the integrated dataset. CCAN is comparable to Seurat v4, SCALEX and online_iNMF (**Fig. 5a**). Although the integration performance of CCAN is slightly inferior to scJoint, its label transferring exhibits higher accuracy in identifying cell types in scATAC-seq data when compared to scJoint and Seurat v4. This is evident in the superior accuracy, F1 score, precision, and recall values of CCAN (**Fig. 5b**). In terms of predicted cell types for scATAC-seq data, CCAN’s results closely align with the golden standard, indicating a high level of accuracy (**Fig. 5c**). This suggests that CCAN achieves a well-balanced integration of data and label transferring, allowing it to accurately predict cell types for scATAC-seq data without compromising the performance of data integration for both modalities.

**Fig. 5.**
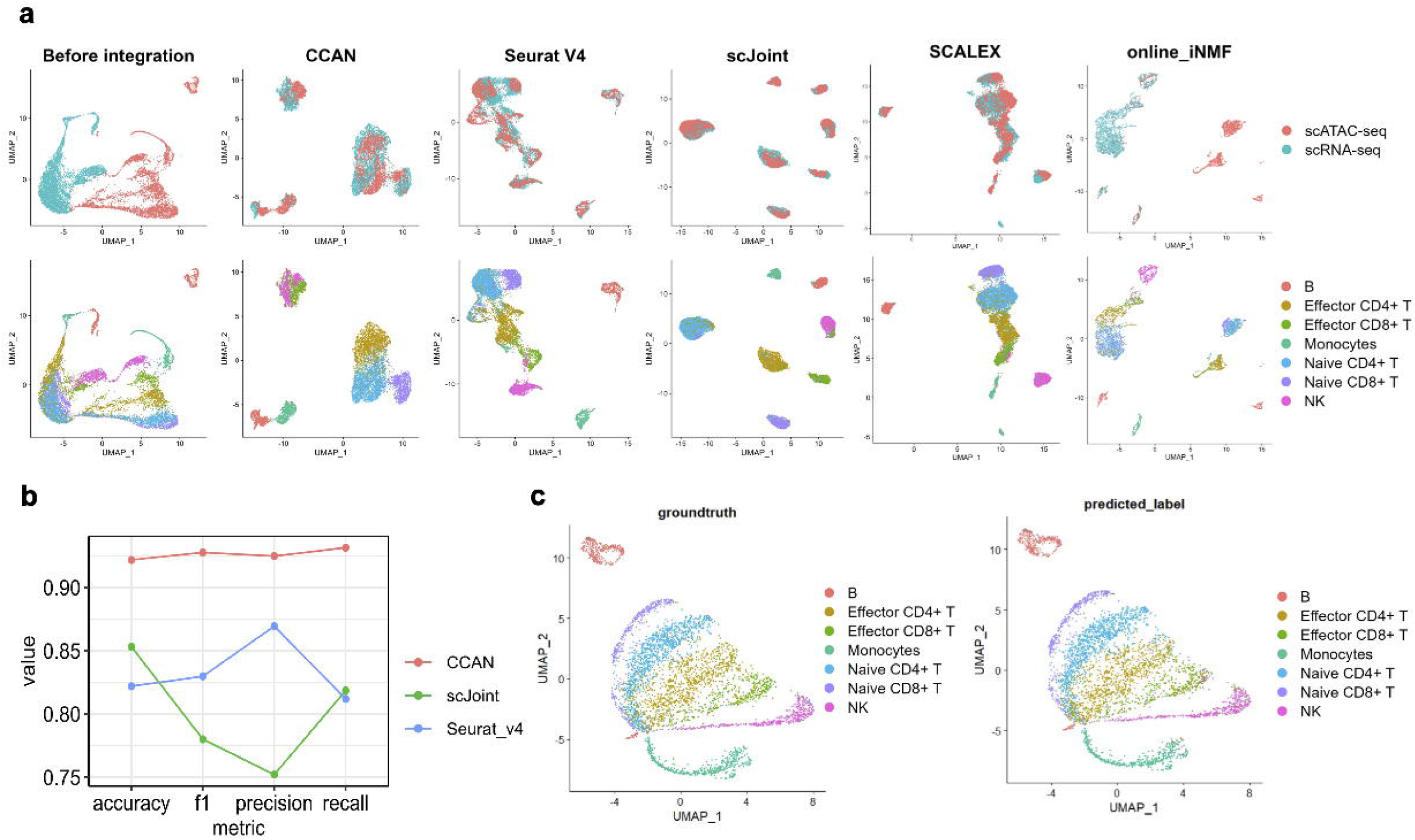
Integration of unpaired datasets and label transferring from scRNA-seq data to scATAC-seq data. **a,** UMAP visualization unpaired **scRNA-seq data and scATAC-seq data** before integration, and integrated embedding after using CCAN, Seurat V4, scJoint, SCALEX, and online_iNMF, respectively. Different colors in the upper panel represent different modalities and different colors in the lower panel represent different cell types. **b,** line plot of four prediction evaluation metrics including accuracy, F1 score, precision and recall based on CCAN, Seurat V4 and scJoint’s prediction results. Different methods were represented using different colors.

### Trajectory inference and pseudotime analysis on the integrated scRNA-seq data and scATAC-seq data

We used human hematopoiesis dataset [49] to evaluate the performance on the integration of differentiated datasets. The human hematopoiesis dataset profiles the chromatin accessibility and gene-expression data of single cells that undergo a differentiation path from hematopoietic stem cells (HSC) dividing into branches. One branch differentiates in to plasmacytoid dendritic cells (pDC), the other goes through common myeloid progenitor (CMP) and differentiates into three different cell types, including megakaryocyte erythroid progenitor (MEP), Monocyte (mono) and granulocyte–monocyte progenitors (GMP) (**Fig. 6a**). The integration of CCAN removed modality-specific variations and mixed all cells in the embedding space (**Fig. 6b**). **Fig. 6c** shows the visualization of inferred trajectories of the hematopoiesis stem cell differentiation process based on the joint embedding of scRNA-seq and scATAC-seq data. Cell are colored with ground-truth cell types. The differentiation trajectory was inferred and smoothed using Slingshot [55]. From the cell type annotation on the inferred cell trajectory of hematopoiesis dataset, we can clearly see three differentiation paths, basically consistent with the differentiation process reported in literature [49] except that pDC is divided into two clusters. In the differentiation of hematopoietic stem cells (HSC), HSC cells go through common myeloid progenitor (CMP) and differentiate into granulocyte–monocyte progenitors (GMP). While GMP cells undergo a series of differentiation steps that include differentiation into earlier progenitor cells and then further differentiation into different types of immune cells such as dendritic cells [56], which is consistent with the trajectory from GMP cells to pDC cells. Pseudotime analysis using monocle3 [57] based on the inferred trajectories confirms the ability of our model in integrating differentiated scRNA-seq data and scATAC-seq data (**Fig. 6d**).

**Fig. 6.**
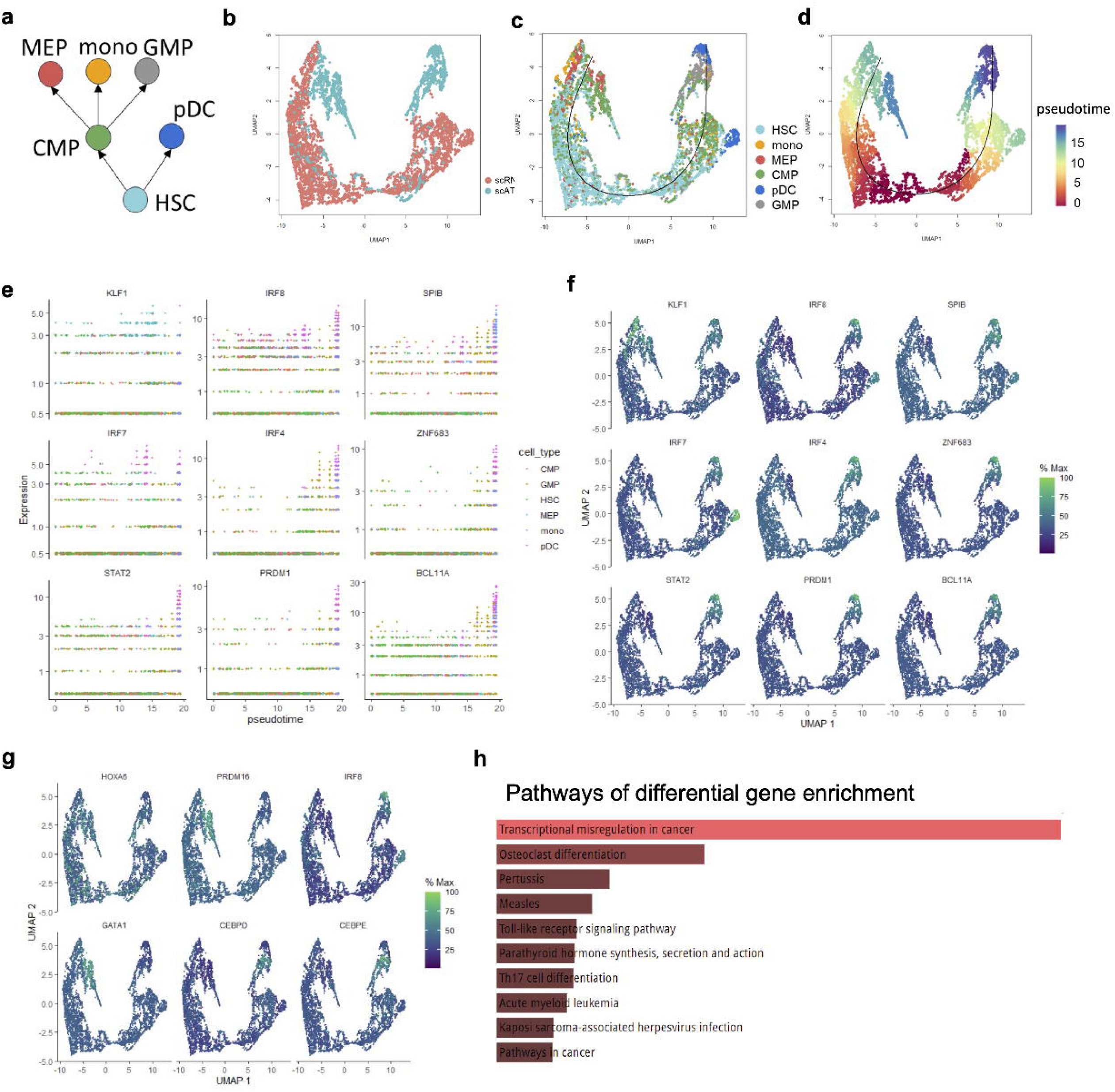
Trajectory inference and pseudotime analysis on the integrated scRNA-seq data and scATAC-seq data. **a,** Reference of human hematopoietic differentiation process from literature. hematopoietic stem cells (HSC) devided into branches, one branch differentiates in to plasmacytoid dendritic cells (pDC), the other goes through common myeloid progenitor (CMP) and differentiates into three different cell types, including megakaryocyte erythroid progenitor (MEP), Monocyte (mono) and granulocyte–monocyte progenitors (GMP). **b,** UMAP visualization of integrated scRNA-seq data and scATAC-seq data colored by different modalities. **c,** UMAP visualization of integrated cells along the inferred trajectory, cells are colored by different cell types. **d,** UMAP visualization of integrated scRNA-seq data and scATAC-seq data colored by differentiation pseudotime inferred by monocle3. **e,** scatter plot of gene expression along the pseodotime of nine highly-correlated genes. **f,** UMAP visualization of integrated scRNA-seq data and scATAC-seq data colored by gene expression of nine highly-correlated genes. **g,** UMAP visualization of integrated scRNA-seq data and scATAC-seq data colored by gene expression of differentially expressed genes identified by DeSeq2. **h,** enriched pathways of differentially expressed genes.

Differential gene analysis can identify differences in gene expression between different cell types during the differentiation process. By calculating the Pearson correlation between gene expression and the inferred cell differentiation pseudotime, we found the nine genes most related to pseudotime, they are *KLF1*, *IRF8*, *SPIB*, *IRF7*, *IRF4*, *ZNF683*, *STAT2*, *PRDM1* and *BLC11A* (**Fig. 6e**). These genes are called transition genes [58]. They are highly expressed in leaf node cells on the cell differentiation trajectory (**Fig. 6f**). According to some published literature studies, the expression of KLF1 is limited to erythrocytes and megakaryocytic-erythroid progenitor cells MEP [59]; *IRF8*, *IRF7*, *IRF4*, *STAT2*, *PRDM1*, and *BLC11A* are all marker genes for plasmacytoid dendritic cells and are highly expressed in pDCs [60–66]. In addition, we also used DESeq2 [67] to identify differentially expressed genes between different cell types. As shown in **Fig. 6g**, *HOXA6*, *PRDM16*, *IRF8*, *GATA1*, *CEBPD*, and *CEBPE* are differentially expressed genes in hematopoietic stem cells, common myeloid progenitor cells, common myeloid progenitors, plasmacytoid dendritic cells, megakaryo-erythroid progenitors, monocytes, and granulocyte-monocyte progenitors, respectively. They are marker genes of different cell type [64, 68–72]. Enrichment analysis based on differential genes across all cell types showed that these differential genes were concentrated in pathways related to hematopoietic stem cell differentiation (**Fig. 6h**). Differential gene analysis further confirmed the effectiveness and accuracy of CCAN in integrating differentiation data with multi-modalities.

## DISCUSSION

Data integration of single-cell multi-omics has enhanced our investigation of cell functions and internal regulatory mechanisms beyond single omics viewpoints. However, single-cell multi-omics integration has numerous challenges, including issues such as omics-variance, sparsity, cell heterogeneity, and confounding factors. The cell cycle confounders in scRNA-seq data inspired us to think about the cell cycle effects in the integration of multi-omics data of cells, especially the integration of scRNA-seq and scATAC-seq data. In this study, we developed CCAN, a cell cycle-aware network for data integration of scRNA-seq and scATAC-seq data and label transferring from scRNA-seq to scATAC-seq. CCAN is based on a domain separation network, which includes periodic activation functions (sine and cosine) in the private autoencoder to simulate and remove cell cycle effects from the integrated single-cell multi-omics data, and projects single-cell data from different platforms or different omics into a common low-dimensional space through shared autoencoder to integrate while maintaining heterogeneity between cell types. The domain adaptive network solves the problem of inconsistent distribution between single-cell data from different platforms or different omics. The class alignment loss is added to the hidden layer of the domain adaptive network to enhance the differences between different cell types in the integrated data. At the same time, by introducing sine and cosine activation functions into the network, the impact of the cell cycle on cell type heterogeneity can be eliminated while effectively integrating single cell multi-omics data, further improving the performance of data integration.

Through comprehensive downstream analyses across diverse data integration scenarios, it has been demonstrated that CCAN (Cell Cycle-Aware Network) possesses the capability to not only mitigate batch effects and cell cycle effects in single-cell RNA sequencing (scRNA-seq) data originating from different platforms but also seamlessly integrate both paired and unpaired scRNA-seq data and single-cell ATAC-seq (scATAC-seq) data. The integration of unpaired scRNA-seq data and scATAC-seq data is particularly noteworthy, as CCAN facilitates accurate cell type prediction for scATAC-seq data by leveraging the transformation of annotation information gleaned from scRNA-seq data. This unique feature enhances the utility of CCAN in deciphering cellular heterogeneity and functional states across diverse data modalities. Furthermore, CCAN exhibits remarkable versatility by not only integrating differentiated data from various modalities but also capturing the intricacies of cell differentiation trajectories. This is exemplified in its ability to characterize the differentiation process from hematopoietic stem cells (HSC) to different branches, providing a holistic understanding of cell fate determination in the context of integrated differentiation data. In essence, CCAN emerges as a powerful tool for unraveling the complexities of cellular dynamics across heterogeneous datasets.

In CCAN, we use a domain separation network that requires the source domain data and the target domain data to have the same number of features. The label transferring in the domain separation network limits CCAN to be used only when the data in the source domain and the target domain have same cell types. While this limitation does not prone to over-correction, which is likely to occur especially when integrating collections of datasets with considerable differences in cellular composition. Currently, there are many methods for analyzing and removing cell cycle effects based on scRNA-seq data, but there are few methods for cell cycle analysis based on scATAC-seq data. Therefore, in this manuscript, the cell cycle effects analysis was focus on the integration of scRNA-seq data from different platforms. When integrating scRNA-seq and scATAC-seq data, we can regard the private-encoder embedding as confounding factors that affecting data integration and cell classification. Besides, we only used single-cell datasets with two modalities, including gene expression profiling from scRNA-seq data and chromatin accessibility from scATAC-seq data. However, with the ongoing advancements in high-throughput single-cell sequencing technologies, the accessibility to analyze various molecular components such as DNA, mRNA, and proteins at a single-cell resolution has expanded significantly. Recognizing the potential for a more nuanced understanding through the integration of diverse omics data types, we envision extending the capabilities of CCAN in future development. The objective is to broaden CCAN’s scope to encompass the integration and comprehensive analysis of a more extensive array of single-cell omics data, thereby facilitating diverse analytical objectives.

## MATERIALS AND METHODS

### Basic structure of domain separation network in CCAN

Domain separation network (DSN) is a specific type of neural network architecture used for domain adaptation, it was designed based on the labeled source-domain data and unlabeled target-domain data [73, 74]. In CCAN, source-domain data is gene expression profiles of scRNA-seq data and target-domain data can be same/different single-cell omics data, such as gene expression of scRNA-seq data from different protocols, chromatin accessibility of scATAC-seq data, etc. In the basic structure of DSN, it contains one shared encoder and two domain-specific private encoders to extract the common and private representations from input data. The common and private components of the same domain should be totally split to make sure the independence of these parts. A shared decoder reconstructs the input domain by cascading the shared and private embeddings. Given a labeled dataset in a source domain and an unlabeled dataset in a target domain, we mainly use cell type classification as the cross-domain task, training on the shared embedding from the source domain that generalizes to the target domain, which requires the high invariance between shared embeddings of source and target domains. To achieve this goal, alignment is considered in the embedding space to eliminate differences between domains. Objectively, DSN is a model that produces a shared representation that is similar for both domains and a private representation that is different and transfers the classification label from source domain to target domain.

### Cell cycle-aware module in CCAN

The cell cycle has been recognized as a confounding factor in the analysis of cell type-dependent processes. In CCAN, to achieve better cell type transfer from the source domain to the target domain, it is necessary not only to account for differences in the shared embeddings between different domains, but also to remove cell cycle effects from the shared representations. Considering the mutual independence between shared and private components in DSN, we introduce a cell cycle-aware module in the private part of DSN to characterize the dynamic process of cell cycle. The high degree of differences between shared and private components can make shared components out of the influence of private components, that is, the prediction of cell types based on shared components can eliminate the effect of cell cycle modeling based on private components. Taking the source data as *X_s_* and the target data as *X_t_* the objective of the cell cycle-aware module is to infer the cell cycle pseudotime *Z_p_* for cells from their corresponding profiles *X_s_* or *X_t_*. DSN is an autoencoder-based network, in the encoder, we use a standard multi-layer perceptron with hyperbolic tangent activation functions in the private part (also as circular part) and selu activation functions in the shared part (also as acyclic part). The encoder of source data *X_s_* is as below

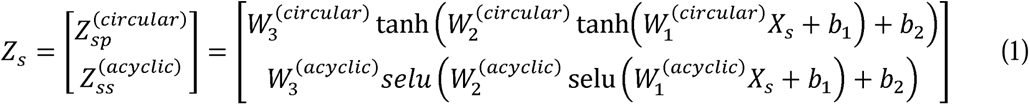

where *W* ’s and *b*’s are the weight matrices and bias vectors of the encoder. *Z_sp_* and *Z_ss_* are private embedding and shared embedding of the source domain. *Z_s_* represents the cascade of *Z_sp_* and *Z_ss_* in the hidden layer. In the decoder, we use cosine and sine as the activation functions in the first layer, followed by two layers performing linear transformations. The reconstruction of source data 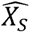 can be represented mathematically as

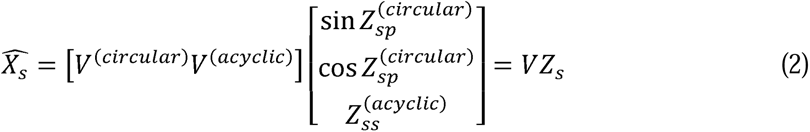

where *V*’s are the weight matrices of the decoder and 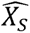 is the reconstructed matrix of source domain generated by the decoder. The target domain has the same autoencoder structure of cell cycle-aware module in the private part as follows.

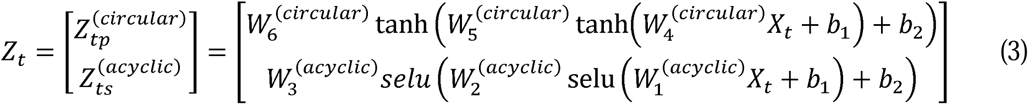

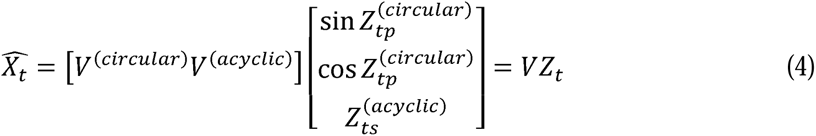

where *Z_tp_* and *Z_ts_* are private embedding and shared embedding of the target domain. *Z_t_* represents the cascade of *Z_tp_* and *Z_ts_* in the hidden layer. 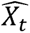 is the reconstructed matrix of target domain.

### Algorithm design and training of CCAN

CCAN uses a domain separation network (DSN) to integrate data from source and target domains and transfer the annotations from source domain to target domain. Shared encoder and private encoders in DSN are three-layer perceptrons to learn circular and acyclic embeddings of both modalities, separately. The shared embedding function projects a high-dimensional profile of each cell to a low-dimensional vector, which distinguishes biological meaningful signals from circular confounding factors (private embedding) and transforms the embeddings of cells from different domains into a similar distribution. In the decoder, we use sine and cosine as the activation functions specific for private embeddings, followed by a two-layer perceptron performing noncircular transformations mapping the embedded data to the original space. CCAN used the labeled transcriptomic profile of scRNA-seq data (source domain) and unlabeled profile from same/different omics data (target domain) as input. Taking scRNA-seq data as source domain and scATAC-seq data as target domain as an example, the training of CCAN has four main steps:

*Step 1: Pretraining of the cell cycle-aware domain separation network.* We used a cell-cycle aware domain separation network to perform joint embedding and modality alignment in a common embedding space through a multi-objective loss. *Firstly,* CCAN trains the source and target autoencoders by minimizing the data reconstruction error.

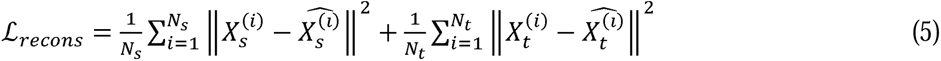

target data, *N_s_* and *N_t_* are number of cells in source and target data, *X_s_* and *X_t_* are input source and target data, 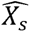 and 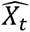 are the decoder-reconstructed data. *Secondly*, it learns shared signals between the scRNA-seq data and scATAC-seq data as well as private signals that are unique to the scRNA-seq data and scATAC-seq data. CCAN applies an orthogonal constrain*L_diff_* to push the features of the shared and the private embeddings apart from each other

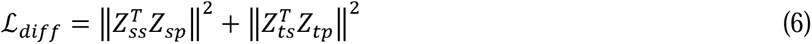

where *Z_ss_* and *Z_ts_* are the embedded source and target data from shared encoder, *Z_sp_* and *Z_tp_* are the embedded data from domain-specific private encoders. The rationale is to disentangle biological signals specific for cell type heterogeneity from cell cycle confounders. *Thirdly*, CCAN regularizes the embeddings of scRNA-seq data and scATAC-seq data to make their distributions to be similar. Aligning the distributions across cells from different domains in the shared embedding space can alleviate the out-of-distribution problem between different modalities. We use Maximum Mean Discrepancy loss *L_MMD_* as the alignment loss *L_align_* to align the distribution of the scRNA-seq data and scATAC-seq data

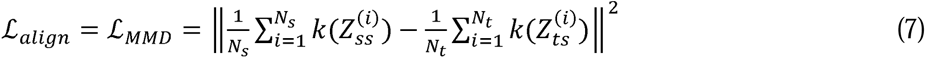

where *k* represents the kernel function. *Finally*, after the unsupervised pretraining of the autoencoder, a supervised cell type classification model can be trained from the aligned shared embedding using the labeled scRNA-seq data, refer as a cross-entropy classification loss *L_class_*

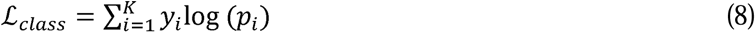

where *y_i_* is ground-truth cell type label of scRNA-seq data and *p_i_* is the predicted probability generated by the cell type classifier. In summary, the total loss in the pretraining of CCAN is

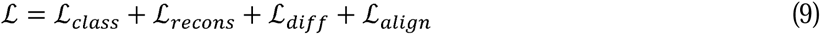

Multiple losses will be given different weights during training.

*Step 2: Label transferring from source domain to target domain.* After pretraining of CCAN, the trained cell type classification network is applied to the unlabeled scATAC-seq data, transferring the cell type information of scRNA-seq data to annotate the scATAC-seq data, we regard it as label transferring. CCAN makes it possible to remove circular confounders while performing accurate annotations for scATAC-seq data. After the label transferring in step 2, we will obtain a pseudo-label of the scATAC-seq data.

*Step 3: Refining CCAN by introducing a cluster alignment loss.* We will improve the joint embedding and label prediction performance using the ground truth of scRNA-seq data and the pseudo-label of the scATAC-seq data. We add a cluster alignment loss *L_ca_* to refine the neural networks in CCAN.

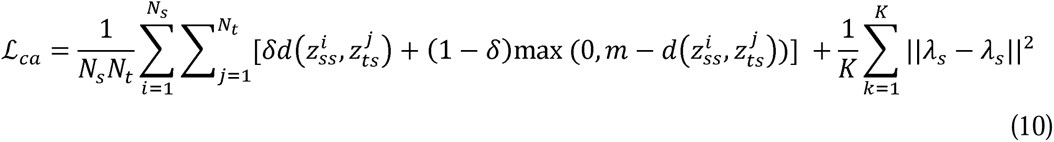

where *m* is the distance threshold, *λ_s_* and *λ_t_* are centroids of embedded source and target data, *K* is the number of cell types. *L_ca_* is a class-conditional loss that forces the features from the same is class to concentrate together and the features from different classes to be separated. In addition, it introduces a conditional feature matching loss to improve the alignment between two domains that aligns the clusters which correspond to the same class but come from different domains. Thus, the updated alignment loss is

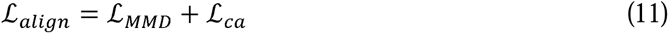

*Step 4: Finalization of CCAN model.* The last step generates the joint profile of two modalities and finalizes the annotation of scATAC-seq data. Besides, more downstream analyses can be operated on the joint embedding profile of scRNA-seq and scATAC-seq data when we cascaded the shared embedding of scRNA-seq and scATAC-seq.

### Balanced mini-batch training for cluster alignment

In the refining step, we introduce a cluster alignment loss in CCAN, which challenges the choice of batch size in neural network training. The batch size selection in the pretraining step is no longer suitable for the refining step to calculate the cluster alignment loss, because it cannot guarantee that each batch size of data can contain all cell types, and may cause extreme imbalance of batch-size data, which will affect the performance of class alignment. To overcome this problem, we applied a balanced mini-batch training in the optimization of the cluster alignment loss that can virtually balance the class ratio of training samples in CCAN. The numbers of samples for each class in a mini-batch are restricted to be the same. This method does not modify or discard cells in the input data, so it can avoid oversampling and under-sampling problems while improving cluster alignment performance, achieving a better integration of single-cell multi-omics data.

#### CCAN parameters

By default, we set the following parameters for CCAN: batch size: 64, the hidden-layer dimensions in the encoder: [512, 256], the hidden-layer dimensions in the classifier: [32, 16], the dimension of latent space: 64, learning rate: 1e-3, number of training epochs: 1000. Parameters are optimized via grid search and may vary based on input data.

### Datasets

CCAN is an effective tool for multitasking. In order to verify the performance of CCAN in different application scenarios, we used several real single-cell datasets, including scRNA-seq data of peripheral blood mononuclear cells from different protocols [33], paired scRNA-seq and scATAC-seq data generated from SNARE-seq technology [27], unpaired scRNA-seq and scATAC-seq data of different cells from the same tissue [48] and single-cell multi-omics data of human hematopoietic differentiation [49, 75] (**Supplementary Table 1**).

#### scRNA-seq data of PBMCs

The scRNA-seq data of PBMCs consisted of two scRNA-seq data, each assayed on the Chromium 10X platform, but using different library construction protocols: 3’ end v1 (3pV1) and 3’ end v2 (3pV2) chemistries. This dataset was pre-processed following the vignette in Seurat (https://satijalab.org/seurat/articles/pbmc3k_tutorial). The 3pV1 scRNA-seq data has 5,356 cells after data pre-processing and the 3pV2 scRNA-seq data has 8,806 cells after data pre-processing.

#### SNARE-seq data

Single-nucleus chromatin accessibility and mRNA expression sequencing (SNARE-seq) is a droplet-based method to simultaneously profile transcriptome and chromatin accessibility in single nucleus. This dataset is regarded as paired single-cell RNA-seq and ATAC-seq data for adult mouse cerebral cortex. SNARE-seq data of adult mouse brain was pre-processed following the vignette in Signac (https://stuartlab.org/signac/1.2.0/articles/snareseq.html). 8,055 cells were used in CCAN after data pre-processing. This dataset is available from Gene Expression Omnibus (GEO) with GEO Series ID GSE126074.

#### Unpaired scRNA-seq and scATAC-seq data

This dataset includes scRNA-seq data from CITE-seq and scATAC-seq data from ASAP-seq. The CITE-seq generates filtered scRNA-seq data based on the condition of mitochondrial reads greater than 10%, number of expressed genes fewer than 500 and total number of UMI fewer than 1,000. For the scATAC-seq data from ASAP-seq, we filtered out cells with a number of peaks more than 100,000 and calculated the gene activity matrices for scATAC-seq data using Signac [54]. After that, 17,668 overlapped genes are selected as the input features of CCAN. This dataset is available from Gene Expression Omnibus (GEO) with GEO Series ID GSE156478.

#### Single-cell differentiated data

This dataset includes scRNA-seq and scATAC-seq of human hematopoietic differentiation. The scATAC-seq and scRNA-seq were performed separately on different cells and there is no paired relationship between cells from the scRNA-seq data and cells from the scATAC-seq data._Six cell types shared by scRNA-seq and scATAC-seq were selected and used in CCAN, they are hematopoietic stem cells (HSC), plasmacytoid dendritic cells (pDC), common myeloid progenitor (CMP), megakaryocyte erythroid progenitor (MEP), Monocyte (mono) and granulocyte–monocyte progenitors (GMP).

### Input format for compared methods

We compared CCAN with several existing integration methods for different integration situations (**Supplementary Table S2**). These methods are developed for different integration types, so they have different format requirements for input data.

#### Harmony, Conos, Scanorama and BBKNN

These methods are designed specific for scRNA-seq data integration. Their required input is scRNA-seq data with gene expression matrix.

#### Seurat V4 and online_iNMF

The scRNA-seq data is gene expression matrix with genes as rows and cells as columns. The scATAC-seq data is gene activity matrix pro-pressed using Signac.

#### scMVP

scMVP takes raw count of scRNA-seq and term frequency–inverse document frequency (TF-IDF) transformed scATAC-seq as input.

#### scAI

The data of scRNA-seq is a matrix with genes as rows and cells as columns. The data of scATAC-seq is a sparse/binary epigenomic profile with regions as rows and cells as columns.

#### scMVAE

The raw count data of scRNA-seq and scATAC data (gene activity format). Row indicates variable (genes and loci), and column indicates sample (cell).

#### scJoint

The input of scJoint consists of gene activity score matrix, calculated from the accessibility peak matrix of scATAC-seq, and gene expression matrix including cell-type labels from scRNA-seq experiments.

#### GLUE

GLUE requires gene expression profile and peak matrix with *AnnData* format as input. In the pre-processing step of scRNA-seq data, highly variable genes were selected using Seurat v3. Then, the data is normalized and scaled, and PCA dimensionality reduction is performed, using 100 principal components by default. For scATAC-seq, GLUE uses LSI dimensionality reduction with a default dimensionality of 100.

#### SCALEX

SCALEX requires sparse matrix of gene expression and gene activity as two batches. Although it is designed for unpaired data, it can be applied to paired data as well via changing the cell names of one batch.

## DATA AVAILABILITY

All data used in this study are publicly available and described in detail in MATERIALS AND METHODS. Two scRNA-seq data of PBMCs were downloaded from https://support.10xgenomics.com/single-cell-gene-expression/datasets/1.1.0/pbmc6k and https://support.10xgenomics.com/single-cell-gene-expression/datasets/2.1.0/pbmc8k. The SNARE-seq data was downloaded from https://www.ncbi.nlm.nih.gov/geo/query/acc.cgi?acc=GSE126074 (GSE126074). The unpaired scRNA-seq and scATAC-seq data were downloaded from https://www.ncbi.nlm.nih.gov/geo/query/acc.cgi?acc=GSE156478 (GSE156478). The single-cell differentiated data was downloaded from https://data.humancellatlas.org/explore/projects/091cf39b-01bc-42e5-9437-f419a66c8a45/project-matrices and https://www.ncbi.nlm.nih.gov/geo/query/acc.cgi?acc=GSE96772 (GSE96772).

## CODE AVAILABILITY

CCAN is an available tool in the GitHub repository (https://github.com/LiuJJ0327/CCAN).

## ACKNOWLEDGEMENTS

We would like to express our gratitude to our colleagues and friends who provided invaluable advice and support throughout the duration of this study.

