## Supplementary Figure for "A Cell Cycle-aware Network for Data Integration and Label Transferring of Single-cell RNA-seq and ATAC-seq"

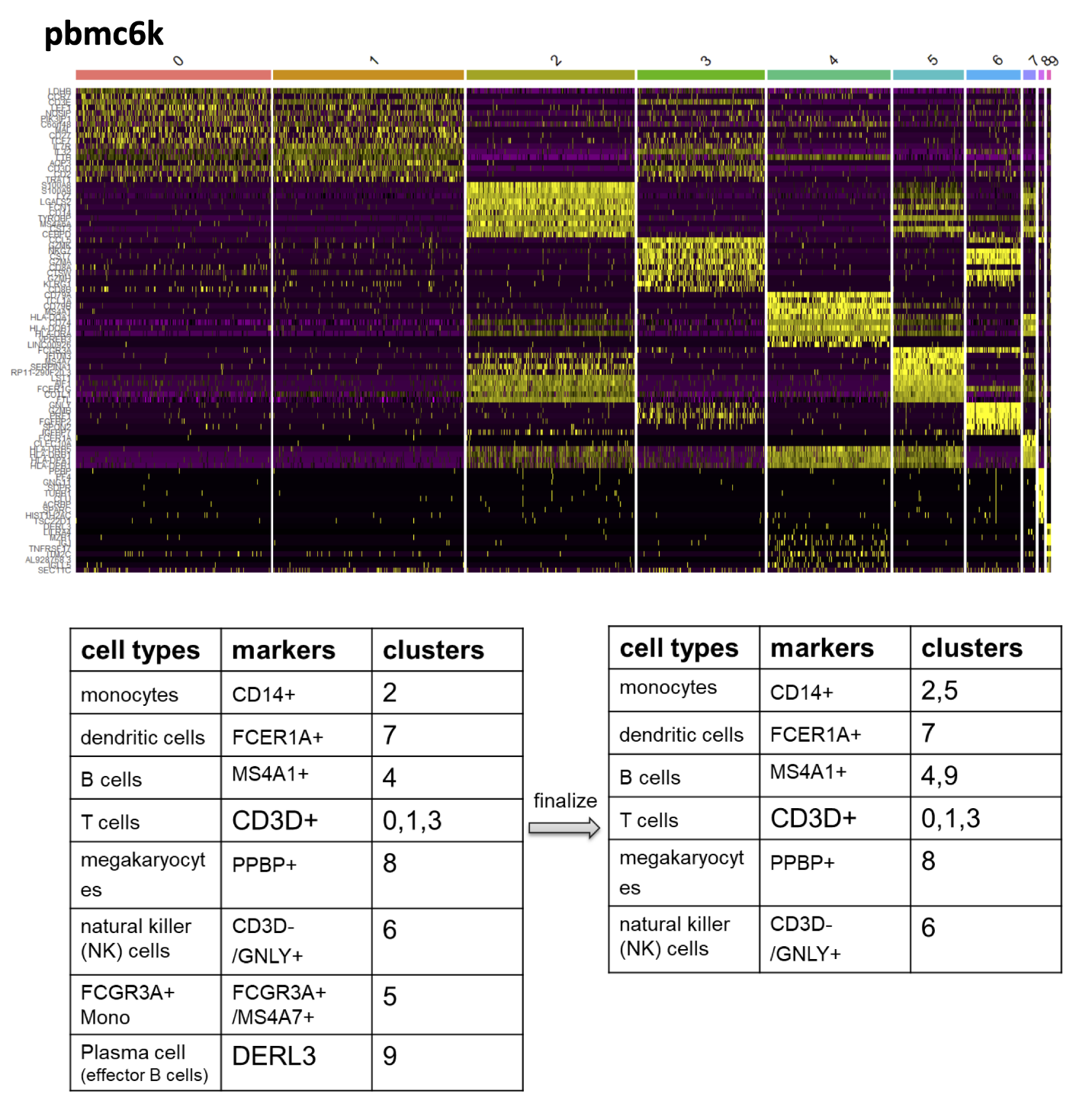

**Supplementary Figure S1.** Cell type annotation of pbmc_6k scRNA-sea data using canonical markers.

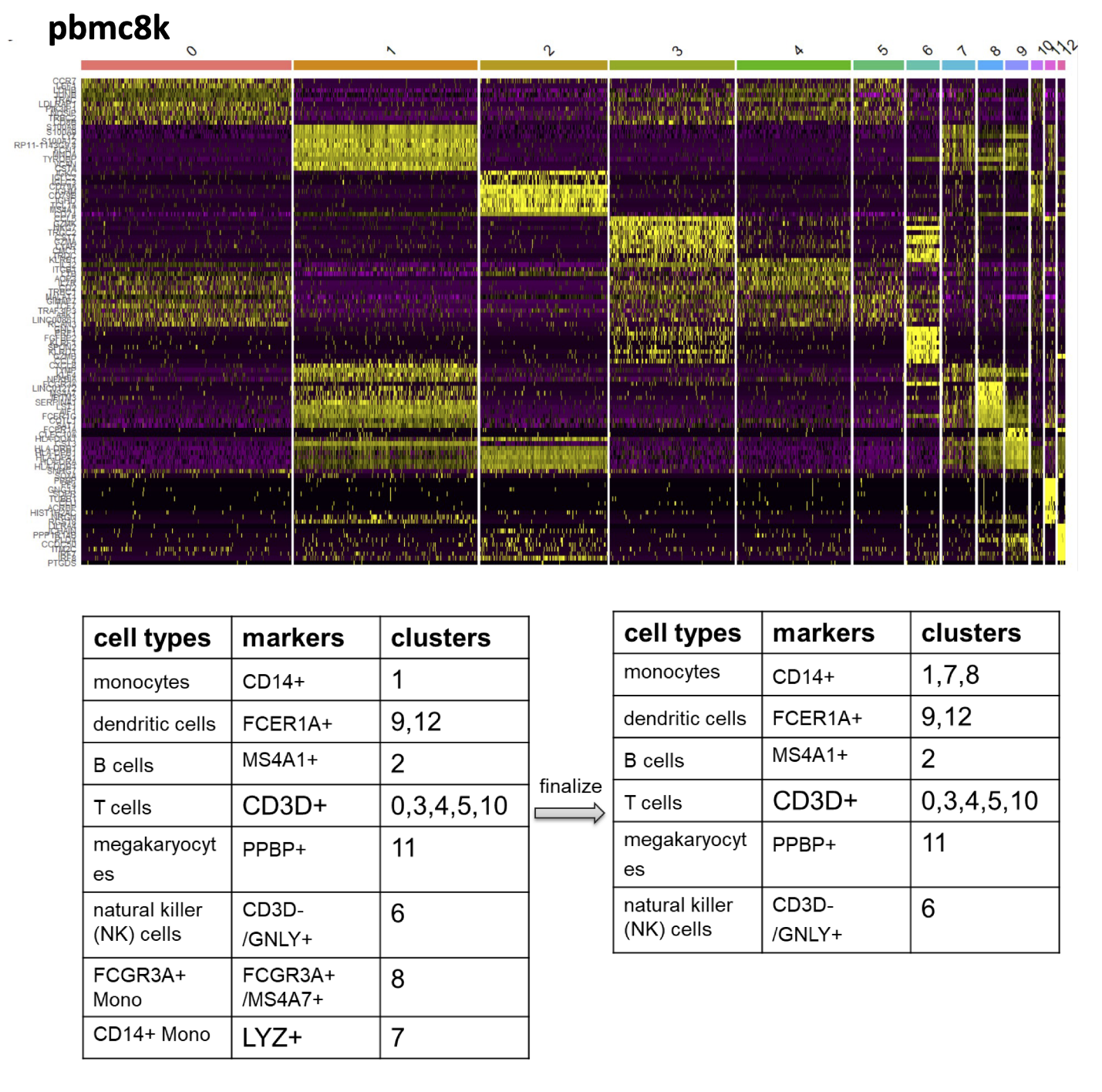

**Supplementary Figure S2.** Cell type annotation of pbmc_8k scRNA-sea data using canonical markers.

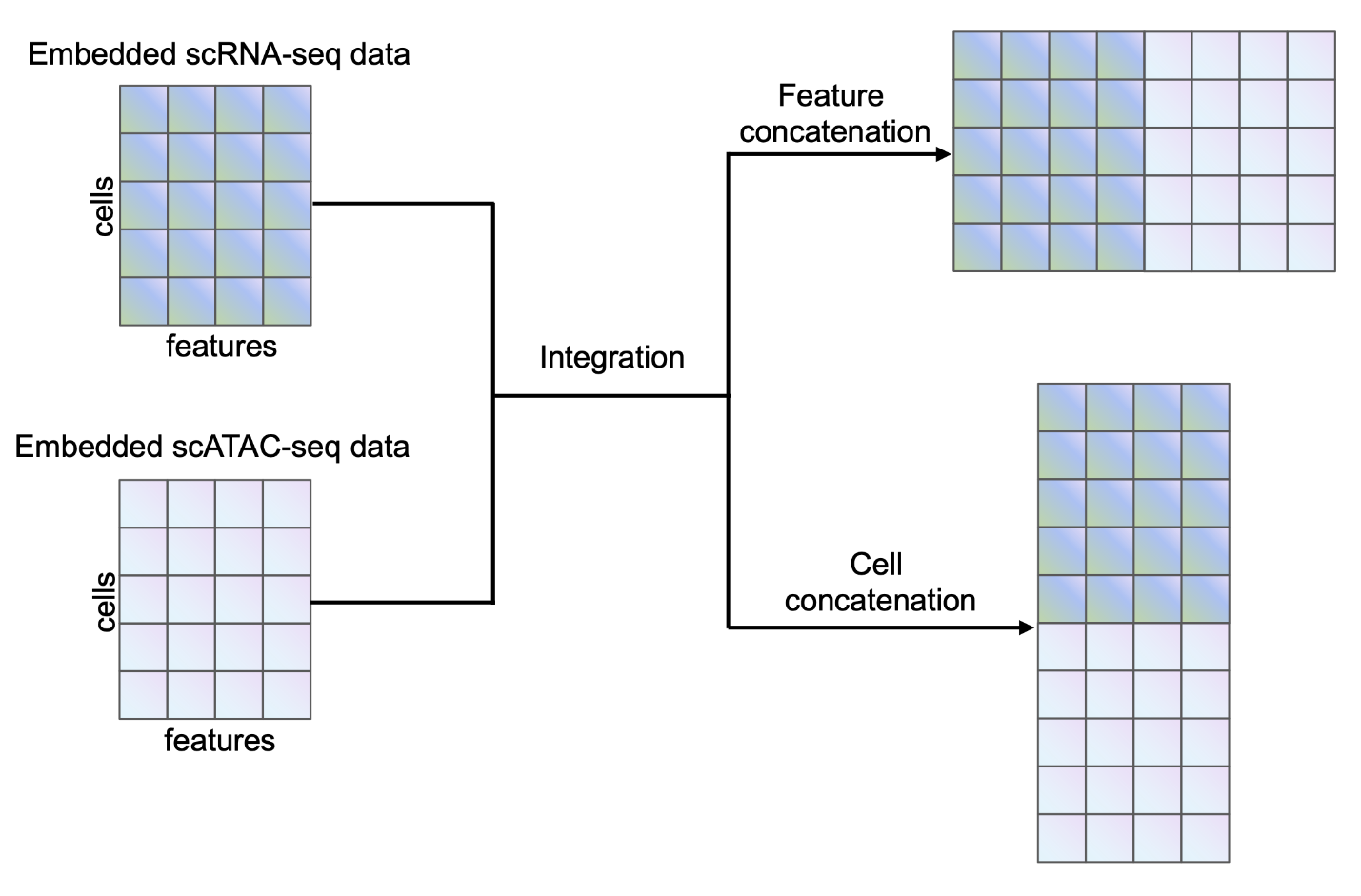

**Supplementary Figure S3.** Illustration of feature concatenation and cell concatenation for paired data when integration.

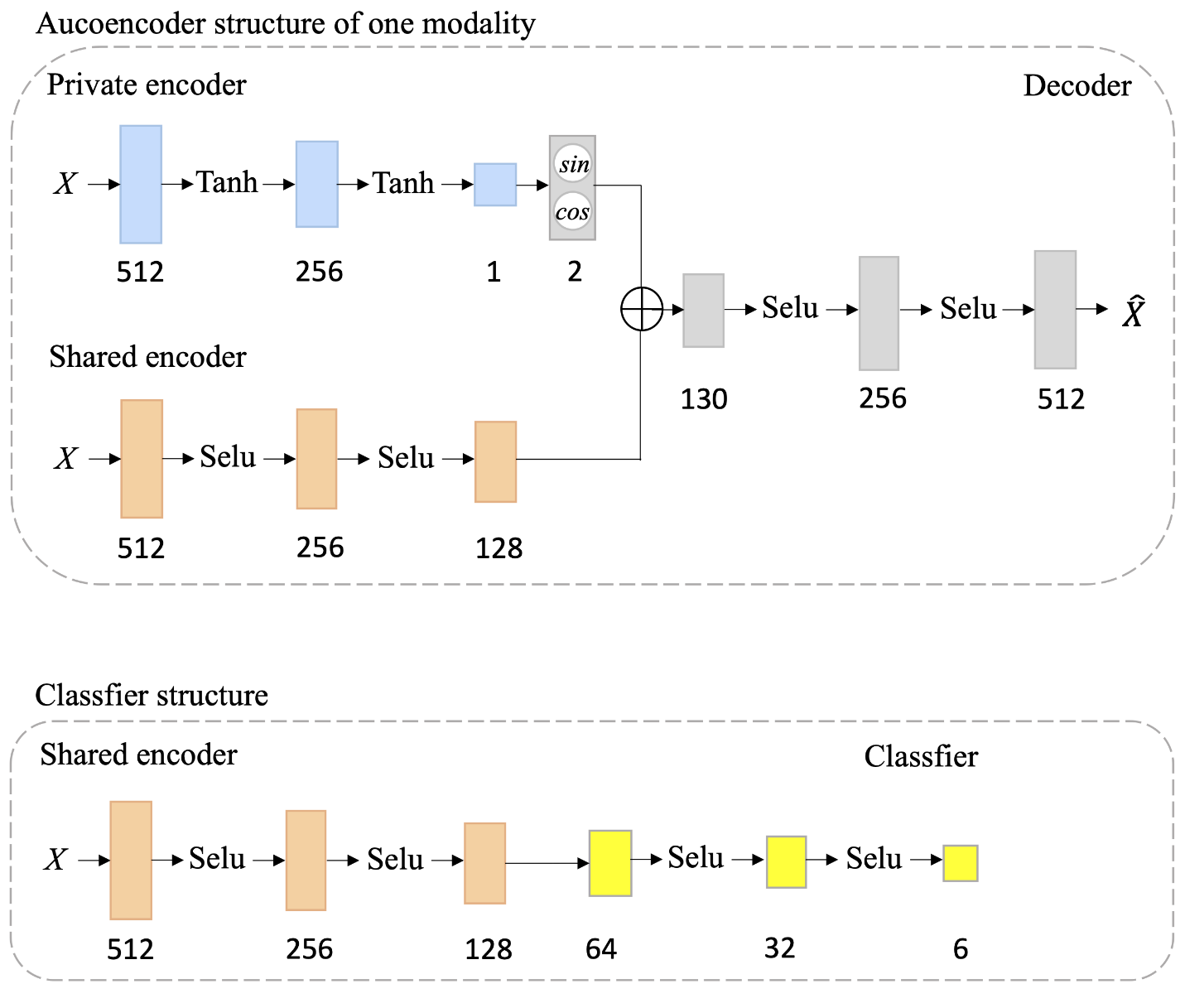

**Supplementary Figure S4.** Network structure of autoencoder and classifier.
